# Hub labeling makes exact distance calculation practical for large human pangenome graphs

**DOI:** 10.64898/2026.11.05.764665

**Authors:** Zia Truong, Adam Novak, Benedict Paten

## Abstract

Pangenome graphs are now a standard reference representation, and tools built on them depend on fast minimum-distance queries between graph positions, for example, when clustering seed matches during read alignment. The baseline distance index, previously used by the vg toolkit, answers such queries based on a decomposition of the graph into nested subgraphs called “snarls”. Within each snarl, the index uses a dense all-pairs distance matrix. Because such a matrix is quadratic in a snarl’s number of children, the large, complex snarls that often occur in many pangenome graphs dominate the overall index size. This means whole-genome graphs can be impractical to index. To mitigate this, the baseline distance index does not fully index the largest snarls, and thus cannot itself answer certain queries involving them.

We replace the dense matrix with a hub labeling for snarls above a size limit. Because hub labeling and associated methods are defined for directed graphs, we introduce a directed representation of a snarl’s bidirected internal “net graph”, whose shortest path lengths are the net graph’s minimum walk lengths. Hub labels are built using pruned landmark labeling, ordered by a contraction hierarchy. On four human chromosome pangenomes, our hub-labeling-based distance index indexes every snarl in 0.9 GB to 3.0 GB at a snarl limit of 500, an estimated 28× to 119× less space than dense distance matrices would need for those snarls with more than 500 children, and supports the 3–18.6% of node-to-node queries that the previous index cannot answer by itself. On the queries both indexes can answer, query time distributions broadly overlap. We explore the snarl limit parameter space and are able to build indexes under 50 GB for four human whole-genome pangenomes in under 6 hours, where we estimate dense matrices could have required multiple terabytes.

## 1 Introduction

Pangenome graphs have become a standard representation of pangenomes for genome-scale sequence analysis. Rather than aligning reads to a single linear reference, projects such as the Human Pangenome Reference Consortium (HPRC) [19] encode many haplotypes in a single graph, capturing population variation directly in the reference structure. The vg toolkit works with these graphs, supporting graph-based read mapping and variant calling at scale [9].

One useful operation on these graphs is that of finding the minimum distance between two graph positions. For example, vg’s Giraffe read mapper relies on such queries to decide which seed matches could belong to the same alignment. For this, it uses a distance index [5], which is based on a decomposition of the graph into nested subgraphs called “snarls” [16]. Within each snarl, the index uses a dense all-pairs distance matrix. This design immediately runs into scaling problems: the per-snarl matrix grows quadratically with snarl size, so a small number of large snarls fill up the index. The **baseline** solution, used in previous versions of vg, has been to use a snarl size limit, dropping all but a few distances for “oversized” snarls, and then to use a runtime graph traversal rather than the distance index to retrieve the missing distances. But this creates a trade-off between index size and the ability to answer queries quickly in complex regions, which is unsatisfying. For some graphs, like those from Pangenome Graph Builder (PGGB) [8], we have been unable to find a point along this trade-off that produces a practical index for read mapping.

One empirically validated approach to exact shortest-path distance in large graphs is “hub labeling”, which answers queries from precomputed per-vertex “labels” and has been shown to scale to graphs with node counts in the millions [13]. To the best of our knowledge, hub labeling has not been applied to bidirected sequence graphs, and no hub labeling algorithm for bidirected graphs was found in the literature.

Here we replace the dense distance matrix of vg’s distance index with a hub labeling, for every snarl above the size limit. (Snarls below this limit still use precomputed distance tables.) To adapt directed graph algorithms to our bidirected graphs, we define a directed representation for the bidirected “net graph” inside each snarl, where shortest path lengths in the directed representation are minimum walk lengths in the net graph. We build labels for the directed representation by pruned landmark labeling [4] ordered by a contraction hierarchy [10], and we adapt vg’s distance index to store these labels. We evaluate the resulting hub-labeling index on four human chromosome pangenomes and four human whole-genome pangenomes, and show that it is practical to use on graphs where the baseline index is impractical.

## 2 Background

### 2.1 Sequence graphs

A **sequence graph** is a bidirected graph (*V, E*) in which each node *v ∈V* carries a DNA **sequence**, whose length we write *l*(*v*). A **variation graph** is a sequence graph representing variation between a collection of related sequences [9].

We follow the conventions of [17] and [5]: we use *orientation* for a direction of traversal of a node and *side* for a left or right endpoint of a node. Thus, each node *v* has two **sides**, a left side *v*^L^ and a right side *v*^R^, and each edge in *E* joins a pair of node sides. (Note that an edge may connect opposite sides of the same node, and may connect one side of a node back to itself.) The **traversal** of *v* from *v*^L^ to *v*^R^ has the sequence of the node; the traversal from *v*^R^ to *v*^L^ has its reverse complement. We call these two traversals the **forward** and **reverse orientations** of *v*, written 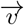 and 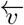.

A **walk** is an alternating sequence of node traversals and edges, where each edge joins the side where the preceding traversal ended to the side where the next one starts. A walk has a length, equal to the total sequence length of all its node traversals. Every walk has a reverse complement walk of the same length, obtained by reversing the order and flipping the traversal orientations.

A **position** is a node traversal together with an offset into its sequence. Here, positions are thought of as being on-base (representing the nucleotides themselves), rather than inter-base (representing the points between, before, or after the nucleotides), and start at 0.

The **minimum distance** between two positions is the smallest number of bases strictly between them, across all possible walks from the first’s traversal to the second’s traversal. When no such walks exist, we say that the minimum distance does not exist, or that it is impossible. The minimum distance between a position and an immediately subsequent position is zero; the minimum distance between a position and itself only exists when there is a cycle in the graph that allows walks to visit the position’s node traversal twice.

### 2.2 Snarl distance index

The **distance index** used in vg [5] answers exact orientation-aware minimum distance queries between positions in a bidirected sequence graph. Indeed, we know of no other index implementation that does so. The index is structured around snarls.

A **snarl** is a subgraph bounded by an ordered pair of traversals that together separate it from the rest of the graph. One or more snarls arranged end to end constitute a **chain**. Snarls can contain nested chains inside them, and those chains contain snarls, forming a tree-shaped structure, the **snarl tree** [5]. The system of snarls and chains captures the intuition that a pangenome is mostly a backbone of shared sequence punctuated by localized bubbles of variation, which themselves are also mostly a few shared sequences with possible nested variation. The snarl tree is, notably, a decomposition of the graph: every node can be hung off of one immediate parent in the tree [16].

Each snarl has a **net graph**: an extension of a sequence graph which describes the snarl’s immediate contents.

Instead of nodes, a net graph has **elements**, which represent the snarl’s children in the snarl tree (nodes and chains) as well as the two nodes bounding the snarl.

Each element in a net graph is assigned a **rank**. In a snarl, rank 0 is the start boundary, rank 1 is the end boundary, and the child nodes and chains are given ranks 2 and above. We also have **root snarls** to represent the root of a snarl tree; these have no boundaries, and their children are given ranks from 0.

A snarl might start or end with the reverse orientation of a node. Additionally, in vg’s version of a snarl decomposition, a child can be marked as reversed in its parent snarl. Elements abstract this away: the right side of the boundary element for the start of the snarl, and the left side for the one at the end of the snarl, always face into the net graph, and the left and right sides of elements inside the snarl are always left and right from the perspective of the containing snarl.

In a sequence graph, nodes can always be traversed left-to-right (or forward) and right-to-left (or reverse), and both traversals are the same length. For elements in a net graph, it might not be possible to traverse from one end to the other (because chains contain snarls, and not all snarls have a walk from one end to the other), so the left-to-right and right-to-left traversals of an element may not exist. An element can also have left-to-left and right-to-right traversals (which can exist when it is possible to turn around in one of the snarls inside a chain), with their own minimum lengths.

Two sides of elements are connected by an edge in the net graph exactly when the corresponding node sides are joined by an edge of the sequence graph.

Walks generalize to net graphs, as sequences interleaving traversals of elements and the edges connecting their endpoints.

The distance index is based around storing all-to-all distance matrices for each snarl’s net graph, and then composing distances in different net graphs to answer questions about distances between positions in the underlying sequence graph [5]. The index is built from leaf to root up the snarl tree hierarchy, so that each parent has access to statistics like the minimum traversal lengths of its children when its own part of the index is being produced. In the baseline index, only snarls below a certain **size limit** are fully indexed, where **size** is defined as the total number of immediate child nodes and chains. **Oversized** snarls (those larger than the size limit) in the baseline index store smaller distance tables that only record distances involving boundary nodes, because the all-to-all distance matrix would be impractically large. When a distance between other elements in an oversized snarl is needed, the query cannot be answered from the index; a shortest-path search needs to be performed at runtime instead.

To extend the distance index to better support large snarls, we need to devise a better way to index minimum walk lengths in their net graphs.

### 2.3 Hub labeling

Hub labeling, also called 2-hop labeling [6], accelerates shortest-path distance queries by precomputing a label for each vertex. It is an exact, rather than an approximate, approach, and has been articulated for undirected and directed graphs. In the directed setting, each vertex *v* of a weighted directed graph receives a **forward label** *L*_*f*_ (*v*) and a **backward label** *L*_*b*_(*v*) [6]. Each is a dictionary of records (*h, δ*) mapping a **hub** vertex *h* to a distance *δ*. For forward labels, *δ* is the distance from *v* to *h*, and for backward labels it is the distance from *h* to *v*.

Labels must satisfy the **2-hop cover property**: for every pair of vertices (*u, v*) where *v* is reachable from *u*, some vertex *h* on some shortest *u → v* path must appear in both *L*_*f*_ (*u*) and *L*_*b*_(*v*).

A query is then a pass over the hubs that the two labels share, adding the two labels’ distances for each hub and taking the minimum. The distance from node *u* to node *v, d*(*u, v*), is given by:

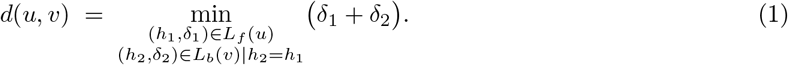

If the two labels don’t share any hubs, the path from *u* to *v* is impossible, which we represent with a distance of *∞*.

Labels can be built by running one Dijkstra traversal in each direction from each vertex and, when the traversal adds a vertex to its “settled” set, adding the source vertex and the Dijkstra traversal’s distance to the settled vertex’s label for that direction. These labels satisfy the 2-hop cover property but are much too large.

To address this, the pruned landmark labeling construction of Akiba et al. [4] adds a **pruning** feature to the Dijkstra traversals: when the traversal rooted at *h* settles a vertex *u* at tentative distance *δ*, the labels built so far are queried for the appropriate path between *h* and *u* using Equation (1). If that query returns a value no greater than *δ*, then *u* is not labeled with *h* and is not expanded in the Dijkstra traversal, because we know the labels already contain some other hub on every shortest path that would contain the *h*-*u* path from the Dijkstra traversal.

This pruning reduces the label size, but since now the Dijkstra traversals depend on each other, it introduces a dependency on the order in which vertices are used to start Dijkstra traversals. In particular, different orderings can result in larger or smaller labels, with different total record counts. Since the cost of a query is bounded by the size of the smaller of the two labels involved, we are interested in finding an ordering that produces small labels [4, 13].

While degree-based orderings [4] and Significant path based Hub Pushing (SHP) [13] have also been used in the literature, here we adopt contraction hierarchies as our ordering heuristic.

### 2.4 Contraction hierarchies

Contraction hierarchies (CH), from the road network literature [10], can provide a vertex ordering for pruned landmark labeling [2]. The method maintains an **importance** score or **priority** for each node, and repeatedly **contracts** the least important remaining vertex, by excluding it from the graph and inserting **shortcut** edges among its neighbors to preserve the lengths of any shortest paths that ran through it. After each contraction, importance scores are updated.

When contracting a node, it isn’t necessary to add a shortcut to replace *every* path from one neighbor to another through that node. When there’s some other equally short path through the rest of the graph, the shortcut edge is not necessary [10]. Avoiding adding unnecessary shortcut edges can speed up the algorithm.

Deciding that a shortcut is unnecessary uses a **witness search**, to find an at-least-as-short path from one endpoint to the other that “witnesses” that the shortcut is unnecessary [10]. Generally one wants to limit the effort expended on witness searches, to avoid more work going into determining that a shortcut edge is unnecessary than would be saved by not having it [10].

While a contraction hierarchy can be used as a shortest-path distance query structure in its own right [10], here we use it just to produce a vertex ordering. The last vertices contracted are, heuristically, the “most important”. They are more likely to lie on many shortest paths [2]. We thus use the reverse of the contraction order as the vertex ordering for building hub labels, so that these more important vertices get added to labels first, creating more opportunities for early pruning during label construction [4].

## 3 Methods

### 3.1 Directed representations of net graphs

To avoid redefining hub labeling, we transform each net graph into a weighted directed graph called the **directed representation**, and compute hub labels on that. See Supplementary A2 for details of the transformation.

### 3.2 Contraction hierarchy implementation

We assign a **hub rank** to each vertex of the directed representation, using a contraction hierarchy (see Section 2.4). For the priority score heuristic, to decide the least important vertex which we then contract, we use the formula of [2].

We adopt the weighting coefficients unchanged from [2], who tuned their function on road networks; we did not retune the coefficients for pangenome graphs.

Contraction proceeds in two phases.

The first phase, though sequential, borrows a technique from parallel contraction hierarchy construction approaches [18] and immediately contracts each vertex whose priority is a local minimum among its un-contracted neighbors, in one pass, such that no two adjacent vertices are both contracted. Because human pangenome graphs tend to be “stringy”, this initial pass cheaply contracts a large fraction of the vertices.

The remaining vertices are then contracted in a lazy priority queue phase [10]. We pop the minimum, recompute its priority, and reinsert it if it no longer has the minimum priority. The last two vertices are ranked without being contracted.

When deciding whether a potential shortcut edge is necessary, our witness search implementation consists of a forward Dijkstra traversal from each in-neighbor of the vertex being contracted. In each phase, the search is bounded by a cap on the number of vertices expanded (250 in phase 1 and 1000 in phase 2), and by a maximum search distance in nodes. This approach is a simplification of the staggered hop limits of [2]. Because the search is limited, we may add unnecessary shortcut edges, which may reduce the quality of our vertex ordering.

Once the order is fixed, the shortcuts are deleted, and only the ranks are kept.

### 3.3 Building labels

Labels are built with the directed variant of pruned landmark labeling [4]. Each vertex is processed as a hub in increasing order of hub rank. Each vertex receives a forward label and a backward label. A record for a hub *h* is added to the forward labels of the vertices that are reachable from the hub with a backward pruned Dijkstra traversal (see Section 2.3). A record is also added to the backward labels of the vertices reachable from the hub with a forward pruned Dijkstra traversal. Each record stores the hub’s rank and the distance found by the traversal.

To avoid seeing 0-length paths from every node back to itself, a hub always appears in its own forward label at distance 0, but not in its own backward label unless a path is actually found back to it.

Because hubs are processed in increasing rank order, label entries are generated already sorted by hub rank. This allows us to easily store the labels as dictionaries from hub rank to distance, represented as sorted lists of records, with lookup performed by binary search. (Since we never need to query labels individually, only intersect them, we do not need to store the mapping from directed representation node ID to hub rank.) The labels for an oversized snarl’s net graph are packed into a flat array and stored in that snarl’s distance index record, which then occupies a region of the distance index’s linear layout. The label array is organized in a columnar fashion, and begins with per-vertex start offsets for the forward labels, then for the backward labels, followed by the hub ranks and the distances of each direction in separate parallel runs. Our distance representation can be up to 32 bits, with the largest signed value reserved to represent unreachability. The whole label array is stored inside the bit-packed distance index vector [5].

### 3.4 Space and time costs

For a snarl with *n* net graph elements, the dense matrix stores a distance for every unordered pair of sides of the snarl’s children, and so occupies *Θ*(*n*^2^) space regardless of the snarl’s internal structure. Our directed representation gives the same snarl 4*n* vertices, and its labeling occupies 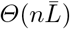 space, where 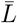 is the mean label size over those vertices. When 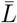 grows sublinearly in *n*, which we hypothesize will be the case in most large snarls from real pangenome graphs, hub labeling will produce an asymptotically smaller index. However, our labeling approach involves labels for all four ports of every net graph element, even those like snarl boundaries where not all ports are used, so hub labelings for very small snarls are likely to be larger than all-to-all distance matrices.

### 3.5 Queries

A query for a distance within an oversized snarl arrives from the distance index query logic in the form of a pair of net graph elements with orientations. (The results of several such queries will be pasted together to answer questions about positions in the underlying sequence graph.) Equation (2) is used to translate the query into directed representation node IDs, which are then translated into directed representation node ranks, which can then be used to query the labels.

To query the hub labels of the directed representation, we use the standard approach of intersecting their labels [6] to compute the result of Equation (1). This is done by iterating over the records of the smaller label and locating each hub rank in the larger label by binary search, an existing approach to the ordered set intersection problem [7]. In general this means that a query should take 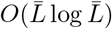 time. Because *both* labels are sorted by hub rank, each search can begin with its lower bound where the previous one ended, speeding it up somewhat. Because the labels are stored in a columnar layout, the binary search operates over memory containing only keys, which will have improved performance relative to searching over interleaved keys and values.

### 3.6 Experimental setup

Indexes were built for two sets of graphs. The first set was four human chromosome pangenomes extracted from the HPRC v1.0 [14] PGGB [8] graph, covering chromosomes 4, 7, 10 and 19. The second set was four human whole-genome pangenomes. Three were HPRC v2.1 “evaluation” graphs [15], built with Minigraph-Cactus [11]. We used the full graph (hprc-v2.1-mc-chm13-eval.full), a version pruned to remove unaligned regions (hprc-v2.1-mc-chm13-eval), and a further frequency-filtered version (hprc-v2.1-mc-chm13-eval.d46). The last graph was the complete HPRC v1.0 PGGB graph that the chromosome graphs above came from (hprc-v1.0-pggb). Table A1 reports graph statistics, and Table A2 reports the size distribution of snarls in each graph.

For the chromosome graphs, we built indexes both with and without hub labeling. The hub labeling indexes were built at snarl size limits 500 and 50000, and the baseline indexes used the vg default of 50 000. Since we did not develop a version of vg that could support both hub labeling and the older baseline method of not indexing oversized snarls, each type of index was built with its own version of vg. Baseline indexes were built with vg v1.75.0 (commit 32d310b, libbdsg 73e74c2) and hub labeling indexes with vg commit a776b6d (based on vg v1.77.0, libbdsg a2e873b). Queries were timed with a small standalone harness linked against each version’s libbdsg. Index construction was single-threaded in all conditions.

For some of the whole-genome graphs, building the indexes without hub labeling was impractical; indeed, we set out to develop the hub labeling method to allow indexing these graphs in practical space and time. Instead of comparing against the baseline indexing approach, we compared a series of snarl size limits (20,500,1000,2000 and 5000) against each other.

Build time is the wall-clock time of one vg index invocation per condition, run on a shared cluster node, measured with GNU `time`. It includes loading the GBZ graph, computing the snarl decomposition, building the index and writing it to disk.

Query time was measured over 500 000 queries per condition, repeated in 3 replicates. We analyzed the median time for each query across the 3 replicates. The set of queries was preselected for each graph, and re-used for all runs on that graph. Each query endpoint was drawn by sampling a node uniformly at random, then an orientation and an offset uniformly within that node. (Note that this biases the query set towards small nodes relative to what we might expect in a real application.) Queries were drawn with the C++ `std::mt19937_64` generator (seed 20261018) using libstdc++’s `uniform_int_distribution`.

The vg distance index is memory-mapped; each index was fully page-faulted from disk into memory before timing any queries, to avoid timing disk I/O.

When querying the baseline distance index implementation, some queries cannot be answered from the index directly, because they would involve distances within the net graphs of oversized snarls. In normal vg operation, these queries would be answered by performing a shortest-path search in the backing sequence graph, but in our experiments we did not load the graph. We excluded these queries from the comparison. This means that, for the chromosome graphs, only the hub labeling condition at snarl size limit 500 is expected to actually exercise hub labeling queries.

#### Compute environment

Jobs were run on any of multiple ostensibly-identical 256-CPU AMD EPYC 7662 nodes, at different times. The nodes reported running Ubuntu 22.04.2, Linux kernel 5.15.0-78, glibc 2.35, and GCC 11.4.0.

### 3.7 Correctness validation

To confirm the correctness of our implementation of hub labeling, we use a collection of unit tests, as well as randomized testing against a Dijkstra traversal that directly finds shortest paths.

## 4 Results

### 4.1 Chromosome pangenomes

#### Index size

The baseline indexes for the chromosome graphs occupy 29.6GB to 444.8GB on disk. Late in our experimental process, we realized that vg v1.75.0 contains a bug that causes its index files to always reserve enough space to hold all the dense snarl distance matrices, even when storing only distances in non-oversized snarls, so these files contain large numbers of trailing zeroes. We were not able to assess the actual *used* space within the baseline indexes by press time. With hub labeling, index sizes range from 0.87GB to 2.99GB at snarl size limit 500, and between 1.88GB and 4.09GB at snarl size limit 50000 (Figure 3, top).

#### Build time

The hub labeling index at limit 500 built faster than the baseline on three of the four graphs (Figure 1, top). The exception is chr7, which builds 54% slower at limit 500 (1331 s vs. 862 s) and 82% slower at limit 50000 (1565s). Its largest snarl, with 210785 children, is the largest in any of the chromosome graphs. Note that build time for the baseline index also includes the time needed to write its large number of trailing zeroes.

**Fig. 1:**
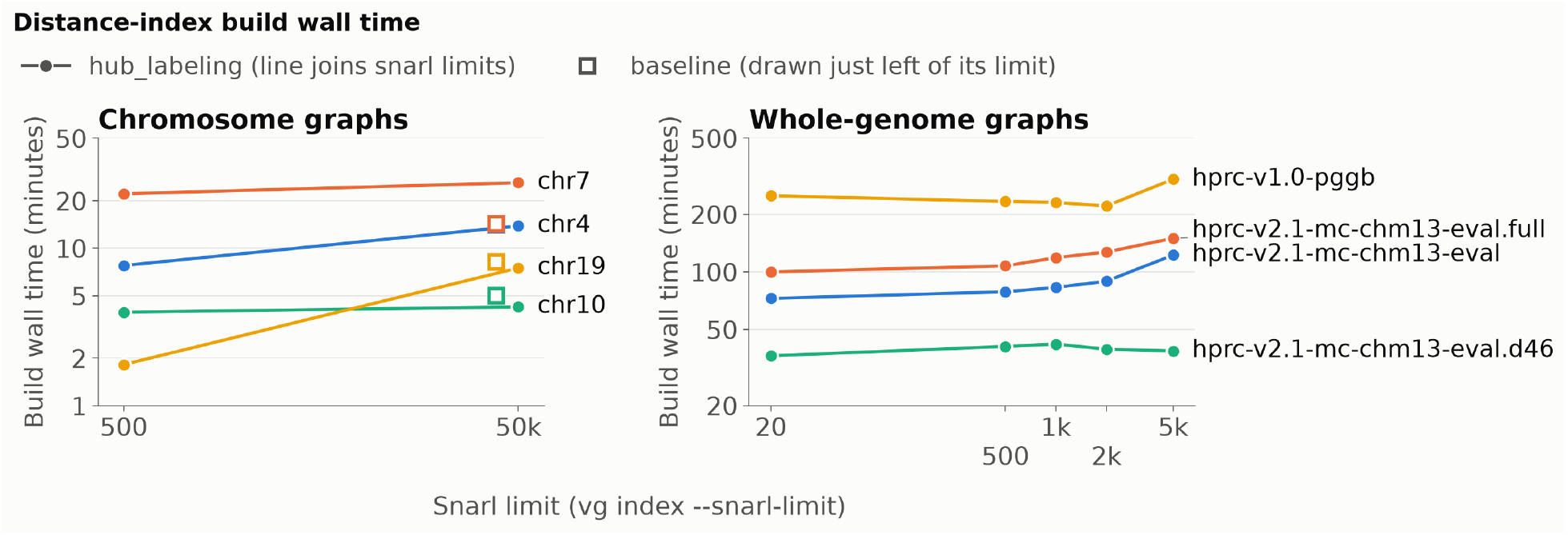
Build time. Top: wall-clock index build time for the four chromosome pangenomes, with hub labeling at snarl size limits 500 and 50 000 (circles joined by a line) and without it at 50 000 (hollow square, drawn slightly left of its limit). Bottom: build time for the four whole-genome HPRC pangenomes at different snarl size limits. All index builds were single-threaded.

#### Query time

On the queries both index types could answer, their median query time distributions broadly overlap (Figure 2, top). All distributions are multimodal. For each graph, median query times differ by less than 10 µs across all three conditions, with trends that differ between chromosomes.

**Fig. 2:**
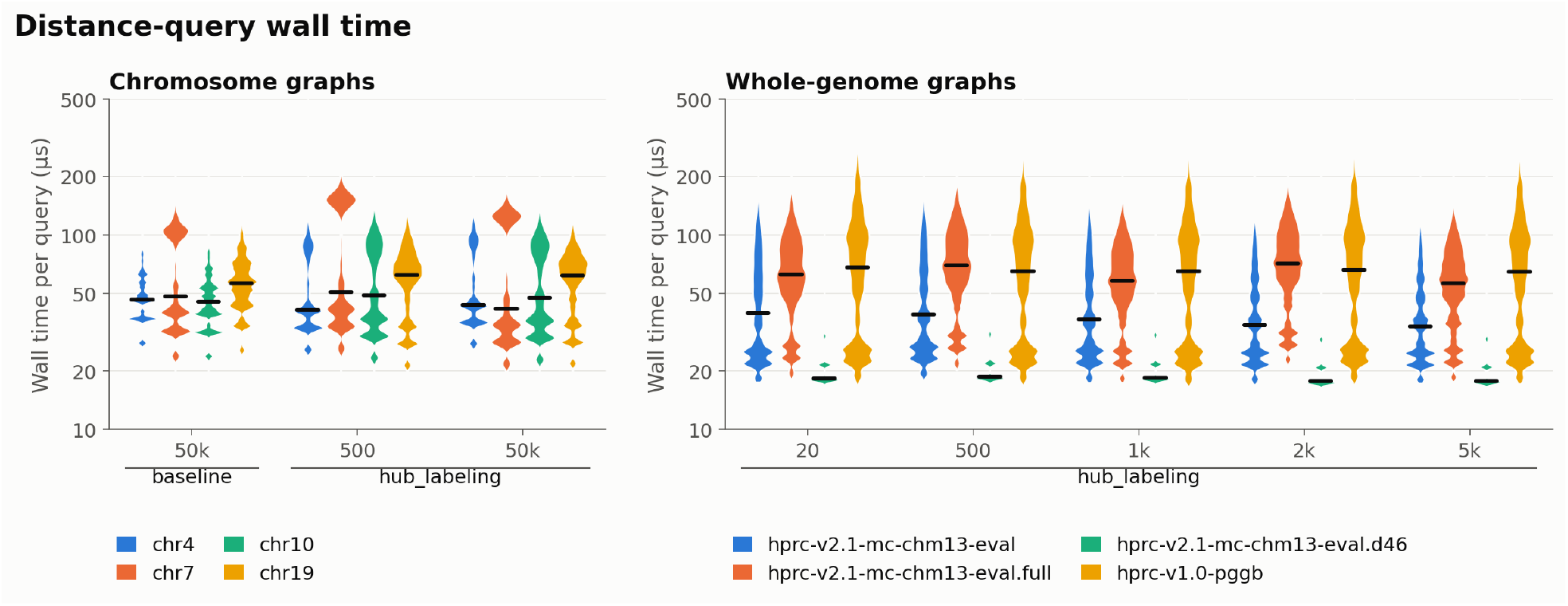
Query time distributions. Each violin shows the distribution of median wall-clock times for each query. Each query’s median time is the median over 3 repeated runs of the same query set. The Y axis is log-scaled. Top: chromosome pangenomes, across the baseline condition and two hub labeling conditions. Note that different chromosomes show different numbers of queries, because queries the baseline condition could not answer were excluded. The baseline index could not answer 3.0% (chr4), 4.7% (chr7), 5.3% (chr10) and 18.6% (chr19) of the 500 000 sampled queries. Bottom: whole-genome pangenomes indexed with hub labeling at snarl size limits 20 to 5000.

**Fig. 3:**
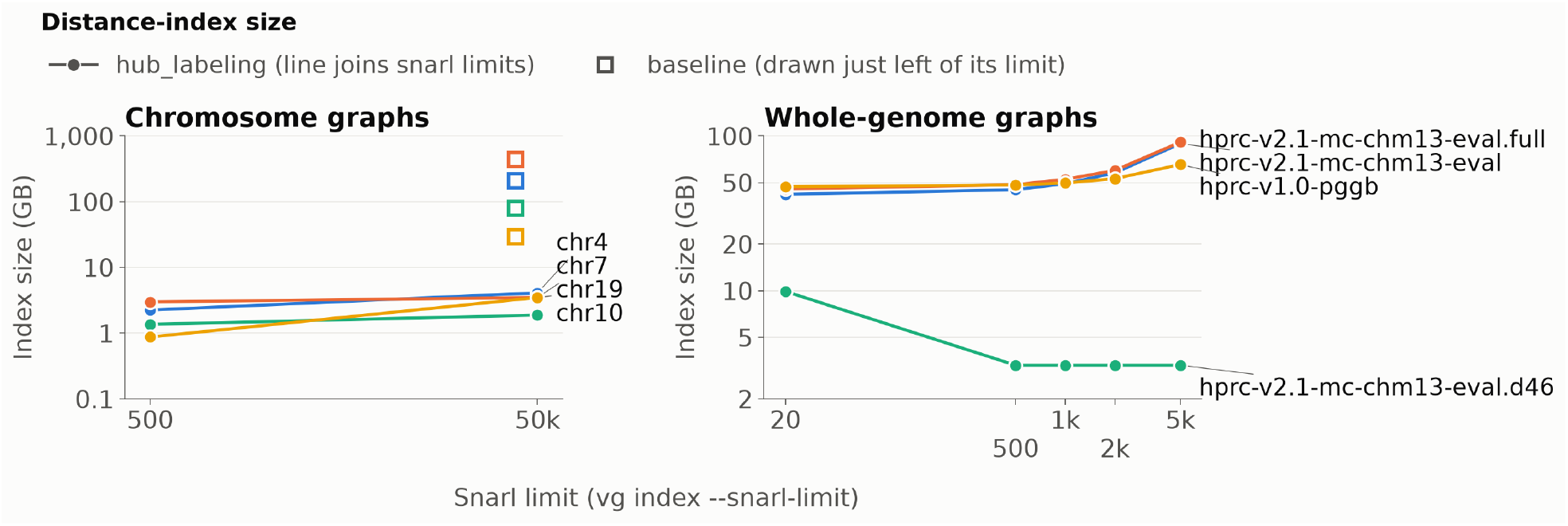
Index size. Size of the serialized index files (decimal GB). Top: sizes for four chromosome graphs across baseline (empty squares) and two hub labeling conditions. Baseline conditions are drawn just to the left of their snarl size limit. Note that baseline sizes reflect the size that would be needed to store dense distance matrices for all snarls, regardless of snarl size limit, due to a bug we found in the baseline version of vg that we are comparing against (see Section 4.1). Bottom: sizes for four whole-genome HPRC pangenomes across hub labeling conditions with various snarl size limits.

We decided to analyze median query times across the three replicates because, when running only one replicate, we noticed that maximum observed query times were oddly high: up to 2.8 ms for the baseline and 28 ms for hub labeling (not shown), about two to three orders of magnitude above the median across queries in a condition. We then changed to using three replicates, and still observed an unusually high maximum in each run, but each replicate had a different slowest query, which took a normal, sub-millisecond amount of time in the other replicates. Operating system statistics suggest that this behavior is not caused by CPU starvation or page faults. Below these maxima is a reproducible slow tail, with high overlap in the slowest 1000 queries across replicates (not shown).

Note that the baseline condition could not answer between 3.0% and 18.6% of queries that involved distances within snarls over its limit of 50000, while the hub labeling conditions could answer all queries regardless of snarl size limit. Queries that could be answered by both implementations always produced identical results. Because of runtime considerations, we did not build the baseline indexes with an unlimited snarl size, which would be necessary to answer the full set of queries. However, we can estimate how large these indexes would be, compared to the hub labeling indexes. Ignoring the low-level bit-packing that the distance index uses, and assuming 4 bytes per entry, the dense matrices for just the snarls of 500 children or larger in each graph would use 25GB (for chr19) to 356GB (for chr7). These estimates track fairly well with the inflated, trailing-zeroes-included sizes of the baseline indexes, and are 28*×* to *×*119 the sizes of the complete hub labeling indexes.

### 4.2 Whole-genome pangenomes

On the whole-genome graphs, where the baseline index cannot be practically built, we swept the snarl size limit through values of 20,500,1000,2000 and 5000 to characterize its effect.

#### Index size

Index size grows with the snarl size limit on three of the four graphs (Figure 3, bottom). For hprc-v2.1-mc-chm13-eval and hprc-v2.1-mc-chm13-eval.full the index roughly doubles between snarl size limits 500 and 5000, from around 45GB to around 90GB; hprc-v1.0-pggb grows more slowly, from around 48GB to around 65GB.

Raising the snarl size limit moves snarls from using hub labeling to using a dense matrix. From snarl size limit 500 to 5000 the number of snarls using hub labeling falls by a factor of between about 5 and 13 (Table A2). On hprc-v2.1-mc-chm13-eval.d46, whose largest snarl has 59 children, the indexes are identical except for at limit 20, where the index grows by about 3 *×*to 9.9GB. This is consistent with hub labeling being larger than dense matrices for small snarls.

#### Build time

Whole-genome builds took between 0.6 h (hprc-v2.1-mc-chm13-eval.d46) and 5.1 h (hprc-v1.0-pggb at limit 5000), single-threaded (Figure 1, bottom), with peak memory of 122GB to 383GB. The hprc-v2.1-mc-chm13-eval.d46 indexes at limits 500 to 5000 are identical, yet their build times range from 2318s to 2516s, a difference of about 8%. This high noise level limits our ability to interpret differences in build time. Broadly, build times seem to rise as fewer snarls are treated as oversized, but on hprc-v1.0-pggb, build times fall between 15020s at limit 20 and 13276s at limit 2000.

#### Query time

Median query time changes only modestly with the snarl size limit (Figure 2, bottom), and not uniformly.

Overall, hub labeling appears at most modestly slower than dense matrix lookup on these graphs. Note that almost all the queries we timed in this experiment (96–97%, depending on the graph) had no connecting walk, often because nodes on two different chromosomes were sampled as endpoints.

## 5 Discussion

We set out to make complete distance indexes practical for pangenome graphs whose complex snarls preclude a dense all-pairs representation of intra-snarl distances. Replacing the per-snarl matrix with a hub labeling achieves our goal: on the chromosome pangenomes it indexes every snarl in at most 3GB, one to two orders of magnitude less than complete dense matrices would need, and it makes whole-genome indexes practical to build. Furthermore, it eliminates situations where the index cannot answer queries and needs to perform path searches in the backing graph.

Our results show that, for hub labeling, lower snarl size limits are not always better for all graphs. On hprc-v2.1-mc-chm13-eval.d46, for example, the index at snarl size limit20is about three times larger than it is at size limit500 (Figure 3, bottom). This is consistent with the space usage analysis in Section 3.4. A hub labeling for a snarl of size *n* costs 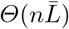 where the matrix it replaces costs *Θ*(*n*^2^), so adopting hub labeling is worthwhile only when the mean label size is well below the net graph size. Dense distance matrices thus still have a place in the vg distance index.

Hub labeling is known to be effective on graphs of “low highway dimension”, a property that formalizes the idea that shortest paths between sufficiently distant vertices tend to pass through a small set of vertices [1]; road networks are the canonical example. Although we have not measured it, we hypothesize that, given what we know about the shape of such graphs, pangenome graphs and the net graphs of the snarls within them would also have low highway dimension, and our present results seem consistent with this. Variation in a variation graph is predominantly local, so a mutation joining two nodes typically joins nodes that are already close in the graph, and long-range shortest paths are thus funneled through shared backbone sequence. Even a complex snarl with no useful decomposition into a few children might still have low highway dimension.

### 5.1 Future work

The contraction hierarchy priority function of [2] was inherited from the road network literature and was not tuned for our graphs. Retuning it could potentially produce smaller labels.

We could also evaluate alternate approaches to constructing a node ordering for hierarchical hub labeling, such as Significant path based Hub Pushing [13]. Other types of labeling could also be explored, such as a directed version of pruned highway labeling [3].

We could also evaluate our implementation on pangenome graphs outside humans, or even cross-species graphs, or specifically on centromeric or other difficult regions. The Minigraph-Cactus graphs we used here for many of our evaluations did not contain variation within centromeres.

We could measure graph statistics like highway dimension [1] or skeleton dimension [12] to try and understand why different pangenome graphs or net graphs are more or less amenable to a hierarchical hub labeling approach.

Finally, having made distance index construction and usage much more practical for large and topologically complex graphs, it is now possible to use these indexes in downstream applications, such as mapping. However, this opens up new challenges downstream and will likely require some further optimization before mappers like those in vg are fast enough to be practically useful for these large and complex graphs.

## Declarations

### Code and data availability

The hub labeling distance index is now available in mainline libbdsg at https://github.com/vgteam/libbdsg and integrated into mainline vg at https://github.com/vgteam/vg. The full genome pangenome graphs (from which the per-chromosome graphs can be extracted) are available from the HPRC at https://github.com/human-pangenomics/hpp_pangenome_resources. Code used for benchmarks can be found at https://github.com/electricEpilith/hublabelBench. Benchmarking results corresponding to this analysis are available at https://github.com/electricEpilith/hublabelBench/blob/1aec230c08d97514329719f1edd1b149d231b53e/results/ (note raw query data is not present due to size).

### Funding

We were in part supported by NIH grants R01HG014490, U01HG010961, U24HG010262, OT2OD026682, UM1HG010971, and U24HG011853. We also received complimentary access to Anthropic’s AI tools during some of the time when the manuscript was being prepared.

### Competing interests

The authors declare no conflicts of interest.

### Author contributions

ZT implemented the hub labeling and contraction hierarchy algorithms. AMN and ZT integrated the algorithms into vg. ZT designed and performed experiments and produced the manuscript and figures. AMN and BP consulted on experimental design and participated in revising the manuscript. BP oversaw the project.

### Use of generative AI

Anthropic’s Claude was used for coding assistance in the implementation described in this paper, for analyzing the existing vg codebase, and for drafting, editing and revising the manuscript. Grammarly was used to reword text and improve grammar. The authors reviewed all generated code and text and take full responsibility for the content of this work.

## Acknowledgements

The authors would like to thank Glenn Hickey for providing graph files, Xian Chang for advice related to snarl distance index code, and Kim Czupil for administering the lab.

## Supplementary Materials

### A1 Graph statistics

Here we give some statistics about the graphs used in our experiments. Table A1 gives summary statistics for the graphs. Table A2 gives information on the distribution of snarl sizes in the graphs. Notably, the hprc-v2.1-mc-chm13-eval.d46 graph has a maximum snarl size of 59, and so, among our tested snarl size limits, only has oversized snarls at snarl size limit 20.

**Table A1:**
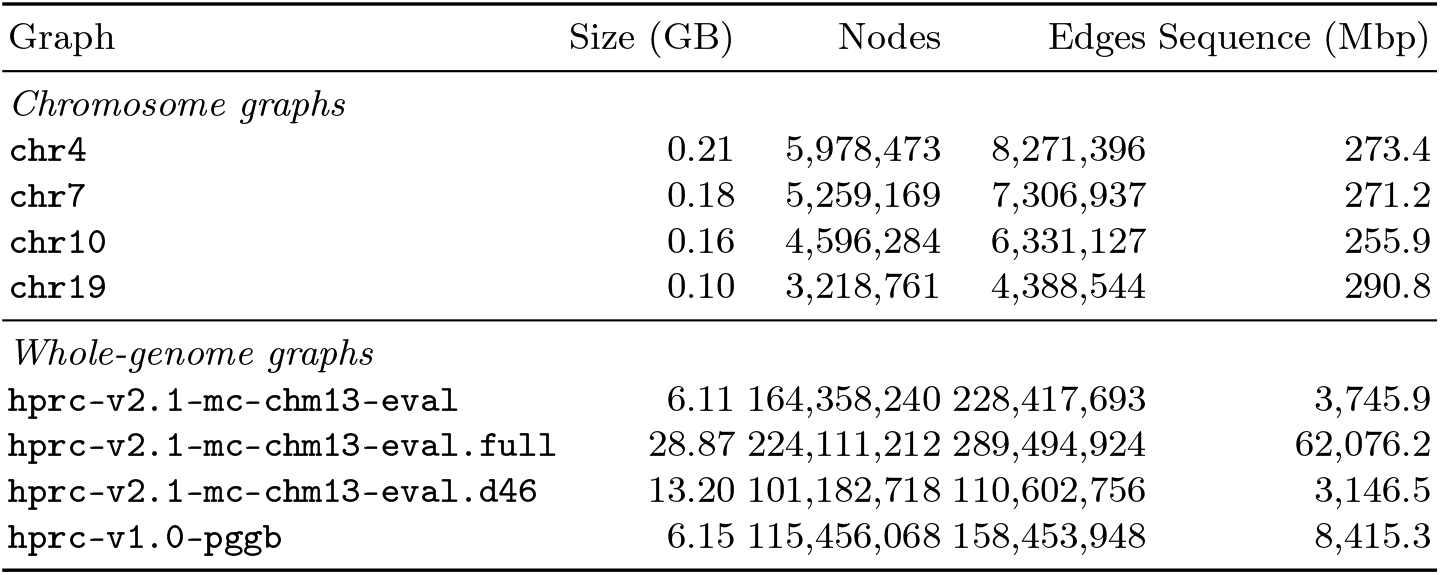
Input graph statistics. Sizes are of the graphs in GBZ format. Nodes, edges and sequence length are from vg stats.

| Graph | Size (GB) | Nodes | Edges | Sequence (Mbp) |
|---|---|---|---|---|
| *Chromosome graphs* |  |  |  |  |
| chr4 | 0.21 | 5,978,473 | 8,271,396 | 273.4 |
| chr7 | 0.18 | 5,259,169 | 7,306,937 | 271.2 |
| chr10 | 0.16 | 4,596,284 | 6,331,127 | 255.9 |
| chr19 | 0.10 | 3,218,761 | 4,388,544 | 290.8 |
| *Whole-genome graphs* |  |  |  |  |
| hprc-v2.1-mc-chm13-eval | 6.11 | 164,358,240 | 228,417,693 | 3,745.9 |
| hprc-v2.1-mc-chm13-eval.full | 28.87 | 224,111,212 | 289,494,924 | 62,076.2 |
| hprc-v2.1-mc-chm13-eval.d46 | 13.20 | 101,182,718 | 110,602,756 | 3,146.5 |
| hprc-v1.0-pggb | 6.15 | 115,456,068 | 158,453,948 | 8,415.3 |

**Table A2:** Snarl size distribution. A snarl’s size is its number of children (child nodes plus child chains). We report total number of snarls (excluding snarls with no children), then count snarls with more than each threshold number of children, which would be deemed oversized at that snarl size limit. We also report the size of the largest snarl.

| Graph | Snarls | >20 | >500 | >1000 | >2000 | >5000 | >50000 | Largest |
|---|---|---|---|---|---|---|---|---|
| *Chromosome graphs* |  |  |  |  |  |  |  |  |
| chr4 | 1,857,464 | 747 | 13 | 7 | 5 | 2 | 1 | 144,895 |
| chr7 | 1,590,933 | 758 | 10 | 5 | 2 | 1 | 1 | 210,785 |
| chr10 | 1,413,582 | 761 | 12 | 4 | 3 | 1 | 1 | 90,222 |
| chr19 | 943,131 | 499 | 11 | 8 | 5 | 2 | 1 | 54,775 |
| *Whole-genome graphs* |  |  |  |  |  |  |  |  |
| hprc-v2.1-mc-chm13-eval | 51,186,896 | 54,351 | 573 | 282 | 137 | 44 | 1 | 828,729 |
| hprc-v2.1-mc-chm13-eval.full | 52,193,498 | 54,894 | 546 | 271 | 146 | 54 | 1 | 839,397 |
| hprc-v2.1-mc-chm13-eval.d46 | 8,714,025 | 167 | 0 | 0 | 0 | 0 | 0 | 59 |
| hprc-v1.0-pggb | 34,523,652 | 15,415 | 255 | 146 | 93 | 53 | 27 | 336,348 |

### A2 Directed representations of net graphs

The hub labeling and contraction hierarchy algorithms are defined for directed graphs.

To avoid redefining them for our net graphs, we transform each net graph into a weighted directed graph whose shortest path lengths are the minimum walk lengths of the net graph, and run the existing directed algorithms on that directed graph. We call this the **directed representation** of the net graph. It is built independently for each oversized snarl. We represent the directed representation using the Boost Graph Library (https://www.boost.org/doc/libs/latest/libs/graph/doc/). The resulting labels are stored as a flat integer array, and the directed representation itself is discarded once a snarl’s labels are computed, so Boost is not involved in querying the index.

Querying requires quickly mapping net graph elements and orientations to directed graph vertices. We achieve this by giving every net graph element a **subgraph** with exactly four vertices in the directed representation, even where fewer would suffice, so that vertex identifiers can be computed in isolation. The four vertices of the element with rank *r* are assigned identifiers 4*r* through 4*r* + 3. The actual ordering of the element ranks should not be very important.

We call each of an element’s four vertices a **port**, defined by a combination of one of the element’s two ends and one of its two orientations. Writing *ρ ∈*{0, 1} for the orientation (0 forward, 1 reverse) and *σ ∈*{0, 1} for whether the port acts as the destination (0) or the source (1) of a query, we use a formula to compute the assigned ID for a port of an element:

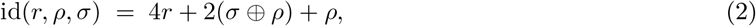

Here, *⊕*denotes bitwise exclusive-or, and the remaining operators denote integer arithmetic.

The second-least-significant bit of the result, which comes from *σ ⊕ ρ*, encodes which end of the element the port sits at (left or right). The least-significant bit, which comes from *ρ*, encodes the orientation. Remember that we define left and right here relative to the net graph’s snarl (see Section 2.2). These two low bits give the **offset** of the port, from 0 to 3.

A port may be either an **in port** (a left forward or right reverse port) or an **out port** (a right forward or left reverse port). Paths representing net graph walks may only arrive in an element’s subgraph through an in port, and may only leave through an out port. Offset 0 is the forward in port and offset 1 the reverse out port, both at the left end, while offset 2 is the forward out port and offset 3 the reverse in port, both at the right end (Figure A4).

**Fig. A.4:**
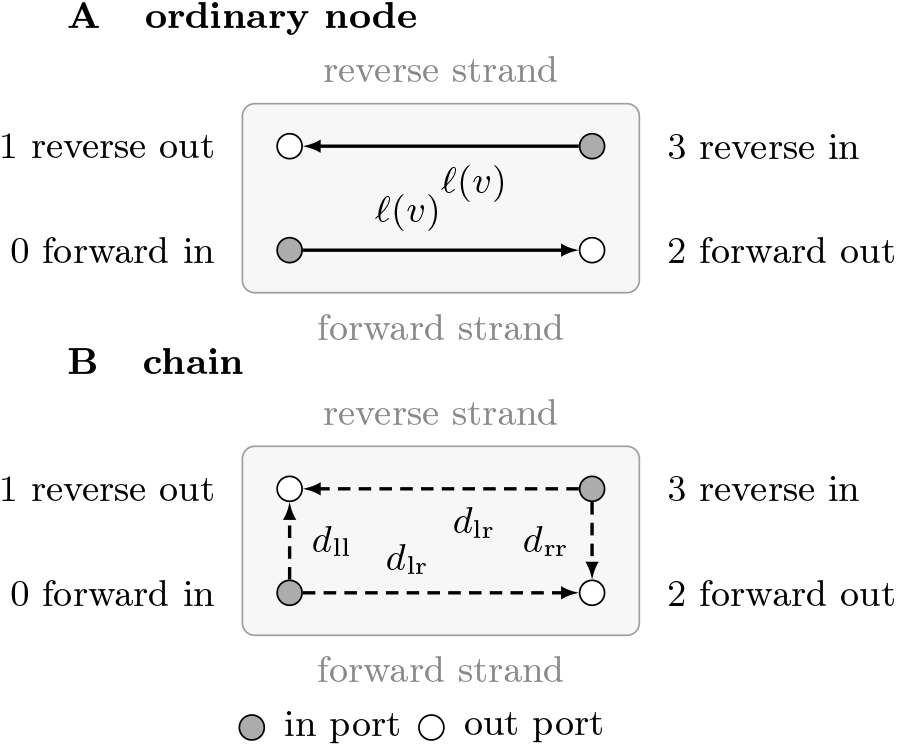
Port layout of net graph elements in the directed representation. An element of rank *r* receives four vertices, one for each combination of end and orientation, at identifiers 4*r* through 4*r* + 3 (Equation (2)). Each element is drawn with its forward strand along the bottom running left to right and its reverse strand along the top running right to left, so offsets 0 and 1 lie at its left end and offsets 2 and 3 at its right end. Filled circles are in ports, at which a path representing a net graph walk can enter the element, and open circles are out ports, at which one can leave. Solid edges are always present. Dashed edges are present only when the corresponding traversals exist. **(A)** An ordinary node gets one straight-through edge per strand, each weighted by the node’s length *l*(*v*). **(B)** A chain gets a straight-through edge on each strand of weight equal to its straight-through, left-to-right (or, equivalently, right-to-left) distance *d*_lr_, if it exists. Where a walk can enter the chain and turn around inside it, it gets an edge across one end of weight equal to its left-to-left distance *d*_ll_ or right-to-right distance *d*_rr_, as appropriate.

We wire the ports in two stages: edges are drawn inside each element’s subgraph to express its internal traversals, and then edges are drawn between elements to express the net graph’s adjacencies. The only nonzero edge weights are for the edges inside element subgraphs.

For a node element, the only traversals are straight through, so we add an edge from the forward in port to the forward out port and an edge from the reverse in port to the reverse out port, each weighted by the node’s length. The two boundary node elements’ subgraphs are built in the same way as for ordinary nodes. For a chain element, we consult the minimum distance information already computed for that child, along with the child’s orientation in the snarl, to obtain its left-to-right, left-to-left, and right-to-right traversal lengths, any of which may be impossible. The left-to-right length is the chain’s minimum length straight through from one end to the other. When that traversal is possible, it yields edges between the in and out ports of the forward strand and between those of the reverse strand, just as for a node. (Because the reverse complement of any walk is also a walk, these two traversals always have the same length and need not be distinguished.)

The remaining two traversals enter the chain and turn around inside it. For chains, the distance index already stores a forward loop distance for the chain’s first node, and a reverse loop distance for the chain’s final node; we combine these with the chain’s orientation in the net graph’s snarl, and the lengths of those first and last nodes, to get left-to-left and right-to-right traversal distances for the chain element, or to know when those traversals are impossible. When the left-to-left traversal is possible, it yields an edge from the forward in port to the reverse out port, and when the right-to-right traversal is possible, it yields an edge from the reverse in port to the forward out port.

Finally, we connect the elements. All edges between elements are zero weight. Each net graph edge joins an unordered pair of (not necessarily distinct) element ends, and produces an unordered pair of (not necessarily distinct) edges in the directed representation: one from the out port of the first end to the in port of the second, and one from the out port of the second to the in port of the first. A net graph edge that joins an end to itself is a **reversing self loop**. At a reversing self loop, a walk can traverse an element in one direction and immediately turn around and traverse it again in the other direction. In this case, the pair of directed representation edges coincide, and only one directed representation edge is created.

The directed representation is designed to have a useful correspondence with the net graph. A directed path from an out port to an in port corresponds to a net graph walk from one element traversal to another, and its total weight is the number of bases strictly between those traversals along the walk. All such directed paths are forced to correspond to net graph walks by construction, because every edge connects one in port and one out port. Edges within elements always arrive at out ports and correspond to routes through the element allowed in the net graph, and edges between elements always arrive at in ports and connect ends of elements that are connected in the net graph.

## Notes

### Competing Interest Statement

The authors have declared no competing interest.

## Bibliography

[1] Abraham, I., Delling, D., Fiat, A., Goldberg, A.V., Werneck, R.F.: Highway dimension and provably efficient shortest path algorithms. J. ACM 63(5) (Dec 2016). https://doi.org/10.1145/2985473, 10.1145/2985473

[2] Abraham, I., Delling, D., Goldberg, A.V., Werneck, R.F.: A hub-based labeling algorithm for shortest paths in road networks. In: Pardalos, P.M., Rebennack, S. (eds.) Experimental Algorithms. pp. 230–241. Springer Berlin Heidelberg, Berlin, Heidelberg (2011)

[3] Akiba, T., Iwata, Y., Kawarabayashi, K.i., Kawata, Y.: Fast shortest-path distance queries on road networks by pruned highway labeling. In: Proceedings of the Meeting on Algorithm Engineering & Experiments. p. 147–154. Society for Industrial and Applied Mathematics, USA (2014)

[4] Akiba, T., Iwata, Y., Yoshida, Y.: Fast exact shortest-path distance queries on large networks by pruned landmark labeling. In: Ross, K.A., Srivastava, D., Papadias, D. (eds.) Proceedings of the ACM SIGMOD International Conference on Management of Data, SIGMOD 2013, New York, NY, USA, June 22-27, 2013. pp. 349–360. ACM, New York, NY, USA (2013). https://doi.org/10.1145/2463676.2465315, 10.1145/2463676.2465315

[5] Chang, X., Eizenga, J., Novak, A.M., Sirén, J., Paten, B.: Distance indexing and seed clustering in sequence graphs. Bioinform. 36(Supplement-1), i146–i153 (2020). https://doi.org/10.1093/BIOINFORMATICS/BTAA446, 10.1093/bioinformatics/btaa446

[6] Delling, D., Goldberg, A.V., Werneck, R.F.: Hub labeling (2-hop labeling). In: Encyclopedia of Algorithms, pp. 932–938 (2016). https://doi.org/10.1007/978-1-4939-2864-4_580, 10.1007/978-1-4939-2864-4_580

[7] Ferragina, P.: Set Intersection. In: Pearls of Algorithm Engineering, pp. 72–81. Cambridge University Press, Cambridge (2023). 10.1017/9781009128933.007, https://www.cambridge.org/core/books/pearls-of-algorithm-engineering/set-intersection/6E6D22DF933E96D87F6EAFC8FB24F688

[8] Garrison, E., Guarracino, A., Heumos, S., Villani, F., Bao, Z., Tattini, L., Hagmann, J., Vorbrugg, S., Marco-Sola, S., Kubica, C., et al.: Building pangenome graphs. Nature Methods 21(11), 2008–2012 (2024)

[9] Garrison, E., Sirén, J., Novak, A.M., Hickey, G., Eizenga, J.M., Dawson, E.T., Jones, W., Garg, S., Markello, C., Lin, M.F., Paten, B., Durbin, R.: Variation graph toolkit improves read mapping by representing genetic variation in the reference. Nature Biotechnology 36(9), 875–879 (Aug 2018). 10.1038/nbt.4227, http://dx.doi.org/10.1038/nbt.4227

[10] Geisberger, R., Sanders, P., Schultes, D., Delling, D.: Contraction hierarchies: Faster and simpler hierarchical routing in road networks. In: McGeoch, C.C. (ed.) Experimental Algorithms. pp. 319–333. Springer Berlin Heidelberg, Berlin, Heidelberg (2008)

[11] Hickey, G., Monlong, J., Ebler, J., Novak, A.M., Eizenga, J.M., Gao, Y., Marschall, T., Li, H., Paten, B.: Pangenome graph construction from genome alignments with minigraph-cactus. Nature biotechnology 42(4), 663–673 (2024)

[12] Kosowski, A., Viennot, L.: Beyond highway dimension: small distance labels using tree skeletons. In: Proceedings of the Twenty-Eighth Annual ACM-SIAM Symposium on Discrete Algorithms. pp. 1462–1478. SIAM (2017)

[13] Li, Y., U, L.h., Yiu, M.L., Kou, N.M.: An experimental study on hub labeling based shortest path algorithms. Proc. VLDB Endow. 11(4), 445–457 (2017). 10.1145/3186728.3164141, http://www.vldb.org/pvldb/vol11/p445-li.pdf

[14] Liao, W.W., Asri, M., Ebler, J., Doerr, D., Haukness, M., Hickey, G., Lu, S., Lucas, J.K., Monlong, J., Abel, H.J., et al.: A draft human pangenome reference. Nature 617(7960), 312–324 (2023)

[15] Lucas, J.K., Hebbar, P., Liao, W.W., Macias-Velasco, J.F., Novak, A.M., Asri, M., Balacco, J.R., Blair, A.P., Bolognini, D., Ebler, J., et al.: Hprc2: A human pangenome reference with near-complete coverage of common genetic variation. bioRxiv (2026). 10.64898/2026.07.21.739710

[16] Paten, B., Eizenga, J.M., Rosen, Y.M., Novak, A.M., Garrison, E., Hickey, G.: Superbubbles, ultrabubbles, and cacti. Journal of Computational Biology 25(7), 649–663 (2018)

[17] Sirén, J., Paten, B., the Human Pangenome Reference Consortium: Gbz-base and gaf-base: Indexed pangenome file formats. bioRxiv (2026). 10.64898/2026.07.10.737775, https://www.biorxiv.org/content/early/2026/07/11/2026.07.10.737775

[18] Vetter, C.: Parallel time-dependent contraction hierarchies. Student Research Project 134 (2009)

[19] Wang, T., Antonacci-Fulton, L., Howe, K., Lawson, H.A., Lucas, J.K., Phillippy, A.M., Popejoy, A.B., Asri, M., Carson, C., Chaisson, M.J.P., Chang, X., Cook-Deegan, R., Felsenfeld, A.L., Fulton, R.S., Garrison, E.P., Garrison, N.A., Graves-Lindsay, T.A., Ji, H., Kenny, E.E., Koenig, B.A., Li, D., Marschall, T., McMichael, J.F., Novak, A.M., Purushotham, D., Schneider, V.A., Schultz, B.I., Smith, M.W., Sofia, H.J., Weissman, T., Flicek, P., Li, H., Miga, K.H., Paten, B., Jarvis, E.D., Hall, I.M., Eichler, E.E., Haussler, D., the Human Pangenome Reference Consortium: The human pangenome project: a global resource to map genomic diversity. Nature 604(7906), 437–446 (Apr 2022). 10.1038/s41586-022-04601-8, http://dx.doi.org/10.1038/s41586-022-04601-8

